# SET8 localization to chromatin flanking DNA damage is dependent on RNF168 ubiquitin ligase

**DOI:** 10.1101/766147

**Authors:** Stanimir Dulev, Sichun Lin, Qingquan Liu, Vildan B. Cetintas, Nizar N. Batada

## Abstract

The DNA damage response (DDR) associated post-translational modifications recruit chromatin remodelers, signaling proteins such as 53BP1 and repair factors to chromatin flanking DNA double strand breaks (DSBs) to promote its repair. Although localization of both RNF168 ubiquitin ligase and SET8 methyltransferase at DSBs is essential for 53BP1’s recruitment to DSBs, it is unclear if they do so via the same pathways. Here we report that RNF168 mediates SET8’s recruitment to DSBs. Depletion of cellular pool of ubiquitin through proteasome inhibition abolished RNF168 and SET8’s localization to DNA damage. Knockdown of RNF8 or RNF168 abolished SET8’s recruitment to DNA damage. Moreover, RNF168 and SET8 form stable complexes *in vivo*. Based on these results we propose a model in which SET8, which despite being a pan-chromatin binding protein, can accumulate several folds at chromatin flanking DSBs through tethering to other proteins that specifically localize to chromatin regions with specific modifications.

## Introduction

The DNA damage response (DDR) pathway is an intricate network which recruits signaling and repair proteins to chromatin flanking DNA double strand breaks (DSBs) to promote its repair via either non-homologous end joining (NHEJ) or homologous recombination (HR). Briefly, DDR starts with ATM-dependent phosphorylation of histone H2AX (γH2AX). MDC1 reads and binds γH2AX to recruit many downstream DDR factors, such as the E3 ubiquitin ligase RNF168 (1, 2).

53BP1 protein contributes to higher order organization of the Igh locus and plays an important role in repair of programmed breaks during VDJ and Class Switch Recombination (CSR) in B-cells (3). Moreover, by preventing resection of DSB ends, it negatively regulates HR. Importantly, in BRCA1-deficient cells, it promotes mutagenic DSB repair and tumorigenesis (4, 5). Two primary histone marks required for 53BP1’s accumulation at DSBs are the histone H4 lysine 20 (H4K20) methylation and H2A lysine 15 (H2AK15) ubiquitylation (6) that are promoted by SET8 (7, 8) and RNF168 (6, 9) respectively. SET8 is the only known H4K20 mono-methyltransferase and formation of H4K20me2 requires H4K20me1 as a substrate. Removal of competing H4K20me2 binding proteins such as L3MBTL1 (10) and JMJD2A (11) are also shown to be required for 53BP1’s accumulation at DSBs. That binding of 53BP1 to nucleosome requires dimerization may be a potential reason for the need for high density of SET8 mediated H4K20me at DSBs. Although both RNF168 ubiquitin ligase and SET8 methyltransferase are necessary for 53BP1’s recruitment to DSBs, it is unclear if these two proteins act via dependent or independent pathways.

In our previous study we showed that SET8 monomethyltransferase localizes to DNA damage and its methylation activity is required during DDR for 53BP1’s binding to chromatin flanking DSBs (7). Inhibition of ATM, HDACs and PARP1 did not abrogate accumulation of SET8 (7) and therefore it is unclear how SET8 accumulates at DSBs. SET8 is known to associate with chromatin via two ways: 1) binding to H4 tail (12, 13) and 2) binding to PCNA (14–17). However, SET8’s binding to DSBs can occur independently of PCNA (7). Here we show that SET8 can accumulate specifically at chromatin flanking DSBs is dependent on RNF168.

## Results

### Ubiquitin is required for SET8 recruitment to DSBs

Previously, we reported that ATM, HDAC or PARP inhibitors do not abolished SET8’s recruitment to DNA damage. As the ubiquitin signaling play an important role in DDR, we tested if inhibition of proteasome, which depletes cellular free pool of ubiquitin, compromises SET8’s recruitment to DNA damage. GFP-SET8 expressing U2OS cells were treated with either DMSO or MG132, a well-known inhibitor of proteasome, and then cells were laser microirradiated (see **Methods**). SET8 accumulated at DNA damage in DMSO treated cells but not in MG132 treated cells (**Fig 1**). Ectopic expression of HA-tagged full-length ubiquitin rescued SET8’s recruitment to DNA damage in MG132 treated cells (**Fig 1**). We conclude that ubiquitin signaling is required for SET8’s localization to DSBs.

**Figure 1.**
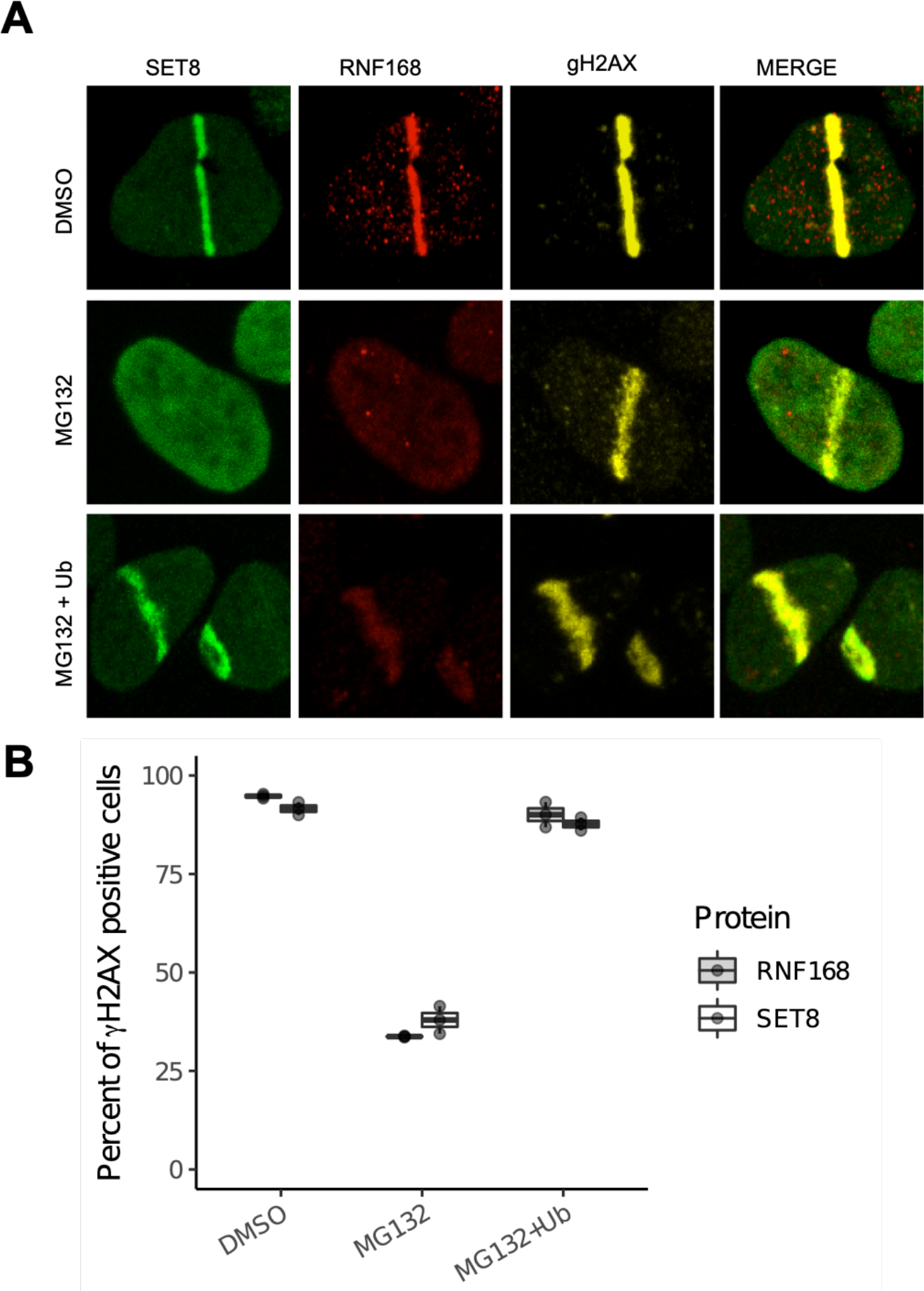
Depletion of cellular ubiquitin pool abrogates SET8’s recruitment to DNA damage. A) Inhibition of proteasome with MG132 abrogates SET8’s recruitment to DNA damage. Ectopic expression of HA-ubiquitin in MG132 treated U2OS cells rescues SET8’s recruitment to DNA damage. B) Quantification of results shown in A. For each of the 3 replicates per condition, over 200 γH2AX positive cells were counted per condition to compute the proportion of γH2AX positive cells that are positive for SET8 or RNF168 (i.e. each dot). Boxplot shows the median of three biological replicates and the box spans the interquartile range.

### RNF168 regulates SET8 recruitment

To identify the potential mediators of ubiquitin modification that play a role in SET8’s recruitment to DNA damage, we depleted E3 ubiquitin ligases RNF8 or RNF168 using siRNAs. While non-targeting control siRNA did not affect formation of γH2AX and recruitment of SET8 and 53BP1 to microirradiation mediated DNA damage, depletion of RNF8 or RNF168 abrogated recruitment of both SET8 and 53BP1 to DNA damage (**Fig 2A, B)**. Thus ubiquitin E3 ligases are important for SET8’s recruitment to DSBs. Since RNF168 is downstream of RNF8, we further interrogated the role of RNF168 in SET8’s recruitment to DNA damage. Endogenous RNF168 were depleted in U2OS cells using siRNA and then cells were transfected with siRNA resistant full-length mCherry-RNF168 plasmid. Expression of mCherry-RNF168, but mCherry alone did not, rescue SET8’s localization at laser-mediated DNA damage (**Fig 2C, D**). These results show that SET8’s recruitment to DNA damage depends on RNF168.

**Figure 2.**
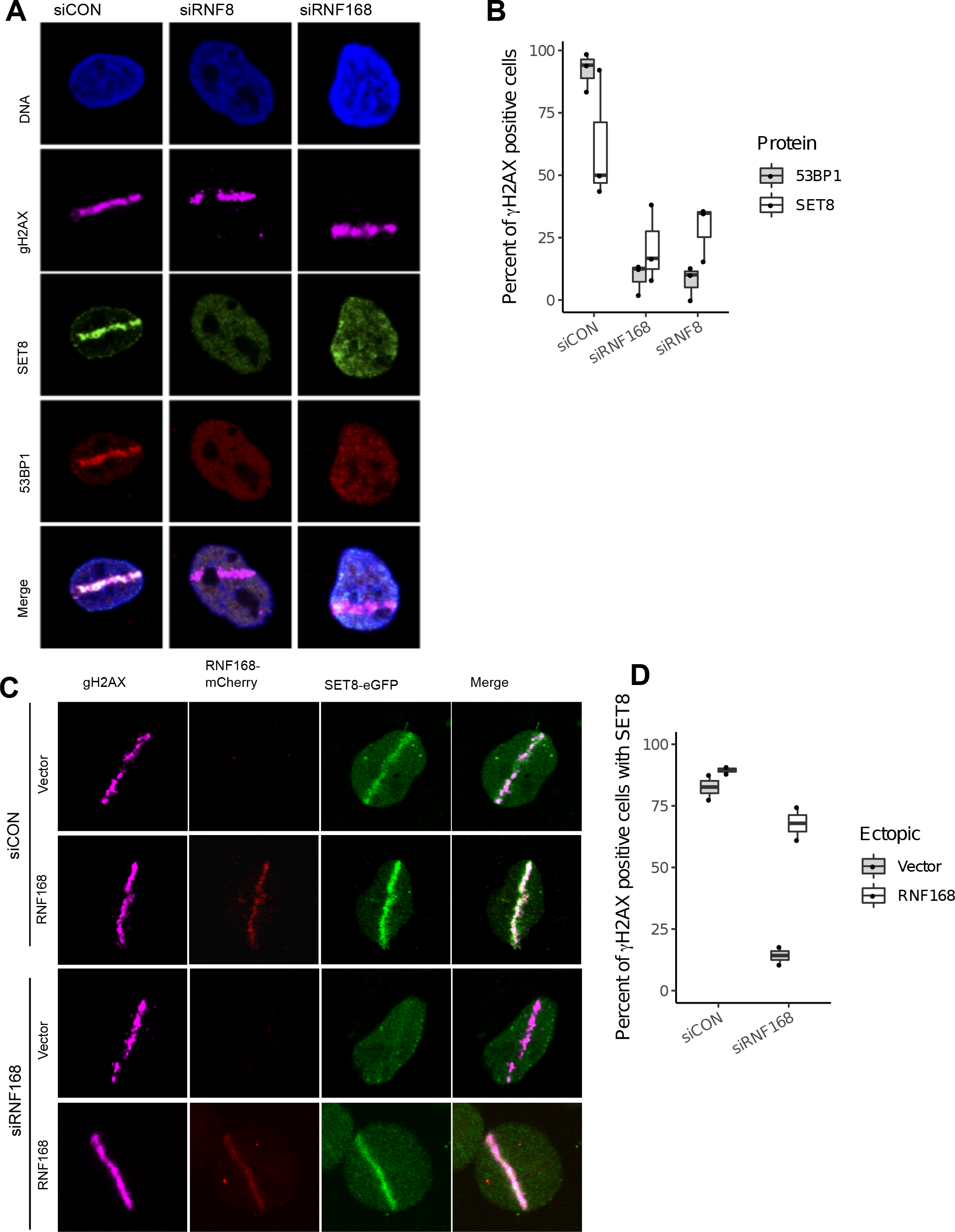
Depletion of RNF8 or RNF168 E3 ligases impairs SET8’s recruitment to DNA damage. **A**) Depletion of RNF8 or RNF168 in U2OS cells abrogates SET8’s recruitment to DNA damage. **B**) Quantification of RNF8 and RNF168 knockdown results shown in A. For each of the 3 replicates per condition, over 200 γH2AX positive cells were counted per condition to compute the proportion of γgH2AX positive cells that are positive for SET8 or 53BP1 RNF168 (i.e. each dot). Boxplot shows the median of three biological replicates and the box spans the interquartile range. **C**) Endogenous RNF168 is depleted via siRNA in U2OS cells. Ectopic expression of empty mCherry fails to rescue SET8 however ectopic expression of siRNA-resistant form of mCherry-RNF168 can rescue SET8’s to DNA damage. **D**) Quantification of RFN168 rescue experiment results shown in C. For each of the 2 replicates per condition, over 100 γH2AX positive cells were counted per condition to compute the proportion of γH2AX positive cells that are positive for SET8 (i.e. each dot). Boxplot shows the median of three biological replicates and the box spans the interquartile range.

### SET8 directly interacts with RNF168

Immunoprecipitation of HeLa cells expressing 3×Flag-RNF168 and GFP-SET8 or vice versa with a GFP-trap system revealed that RNF168 an SET8 are part of the same protein complex *in vivo* (**Fig 3A**). We used a potent DNAase (benzonase) during immunoprecipitation, to rule out the possibility that the observed interaction is mediated through DNA. The interaction was present in both irradiated and irradiated cells; others have also observed interaction of other DNA damage proteins in the absence of irradiation. Moreover, the interaction appeared to highly stable as the interaction was seen even under 400 mM salt concentration. To corroborate the interaction between SET8 and RNF168 *in vivo*, we employed the LacO/LacR system to visualize their co-localization on chromatin (**Fig 3B**). HeLa cells with integrations of LacO arrays were transfected with the LacR-mCherry-RNF168 and SET8-GFP plasmids and visualized under confocal microscope. Co-localization of RNF168 and SET8 was clearly observed (**Fig 3C**). We conclude that a certain nuclear pool of SET8 interacts with RNF168 constitutively before and during DNA damage.

**Figure 3.**
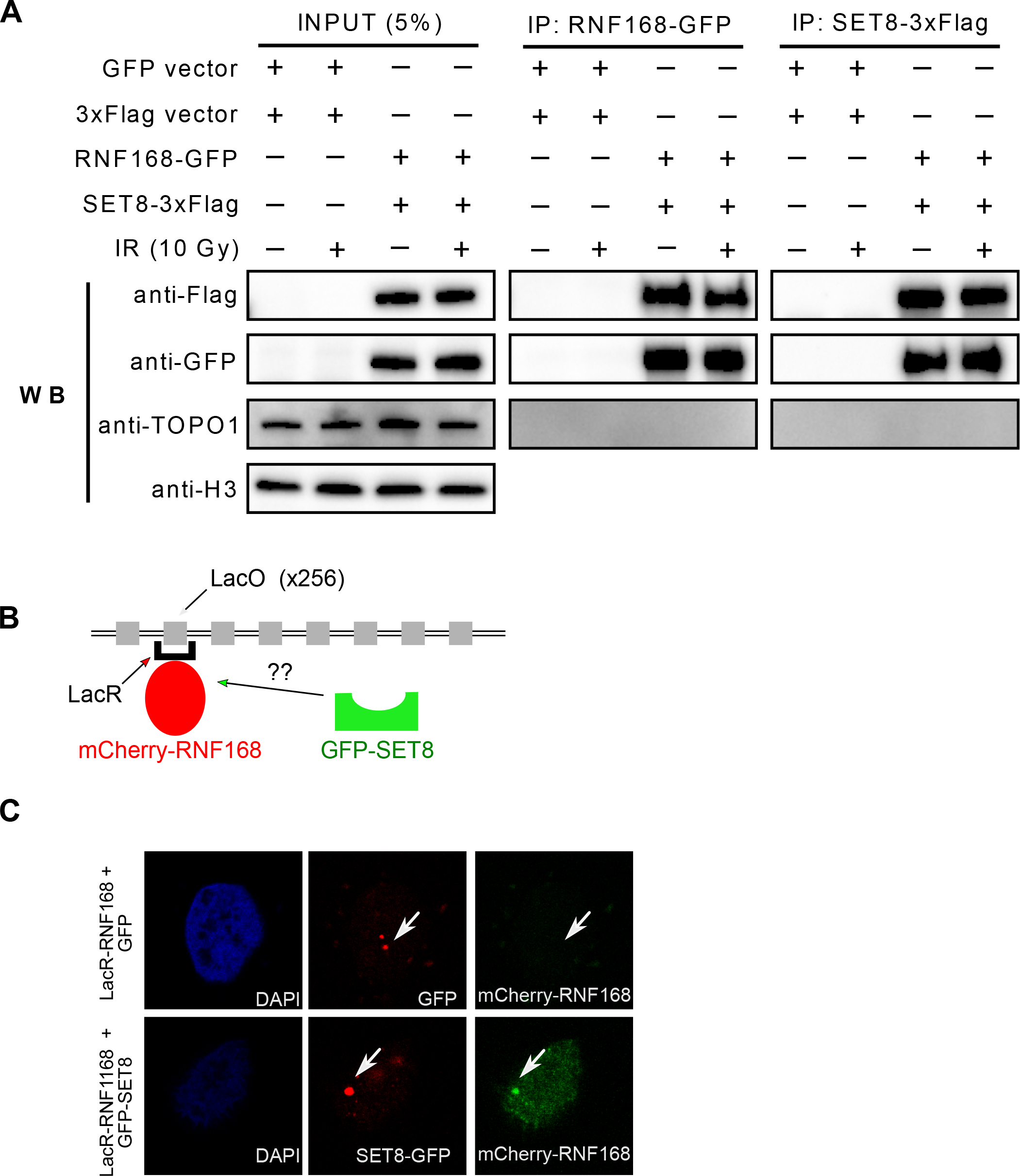
RNF168 and SET8 interact ***in vivo***. A) HeLa cells were transfected with 3×Flag-SET8 and GFP-RNF168 and IP was performed with anti-FLAG beads (lanes 1) or anti-GFP (lane 2). Stable interaction is observed in irradiated (IR) and unirradiated cells. B) Illustration of the LacO array, LacR/ mCherry-RNF168 and GFP-SET8 constructs. C) Assessment of SET8 and RNF168 co-localization *in vivo* in HeLa LacO cells transfected with LacR-tagged mCherry-RNF168 + GFP as a control group and LacR-tagged mCherry-RNF168 + GFP-SET8. Arrows mark the local accumulation of tethered LacR repressor fusions to the LacO operator and the potential colocalization with SET8 and RNF168.

## Discussion

Chromatin modification and remodeling is essential for proper repair of DSBs in human cells. Previously, we demonstrated that the SET8’s methyltransferase activity and localization at DSBs is necessary for 53BP1’s recruitment to DSBs. Here we show that SET8’s role in DDR requires RNF168. In particular, we have shown that depletion of free pool of ubiquitin (**Fig 1**) or RNF8 or RNF168 (**Fig 2**) both abrogate SET8’s recruitment to microirradiation mediated DNA damage. Moreover, we have ruled out siRNA-off target effect by expressing siRNA-resistant form of full length RNF168 to rescue the recruitment of SET8 to DNA damage. Lastly, we have shown via independent methods that RNF168 and SET8 are part of a complex in the cells (**Fig 3**). Thus, we have demonstrated that SET8’s recruitment to DSBs depends on the canonical DDR pathway.

SET8 can localize to chromatin by binding to unmodified histone H4 tails. There are approximately 1 million nucleosomes in the human genome; thus SET8s specific accumulation at chromatin flanking DNA damage (7) would require either additional epitope that would increase affinity of SET8 for nucleosomes near DNA damage by a million fold. Although further experiments are required to properly tease out the exact mechanism via which RNF168 promotes SET8’s recruitment to DSBs, one possibility we favour is that SET8’s localization to DSBs occurs via anchoring on to RNF168. According to this model, the need for ubiquitin for SET8’s localization to DNA damage (**Fig 1**) arises due to the need for RNF168’s recruitment to DNA damage. Simultaneous recruitment of RNF168 and SET8 would promote rapid recruitment of 53BP1 to DSBs as the two post-translational modifications required for 53BP1’s localization at DSBs (i.e. histone H4-K20me1/2 and histone H2A-K13/K15ub) occur simultaneously.

## Materials and methods

### Cell culture

Human U2OS osteosarcoma cells were grown in high glucose DMEM with GlutaMAX or McCoy’s medium containing 10% fetal bovine serum (FBS) in 37°C under humidified atmosphere with 5% CO_2_. Hela cells were cultivated in DMEM supplemented with 10% FBS.

Proteasome inhibitor MG132 was obtained from Calbiochem (San Diego, CA) and dissolved in DMSO. For the MG132 treatment, U2OS cells were seeded on 6 well plates at a density of 1×10^5^ and either treated with the DMSO or MG132 (final concentration 5 μM), and incubated for 30 minutes. For the rescue of MG132 depleted cells, cells were transfected with his-tagged ubiquitin construct using Lipofectamine and incubated for 48h. After the incubation periods, all cell groups were washed with culture medium and micro irradiated before immunofluorescence.

### Plasmids

*H*is-tagged ubiquitin construct was obtained from Addgene (plasmid: #18712). *siRNA*#*1 resistant RNF168*-mCherry *plasmid and RNF168 siRNAs were gift from Jiri Lucas (1). SET8 was subcloned into pcDNA5/frt/to-GFP vector (Invitrogen)*.

### Laser Microirradiation

U2OS cells were seeded on 25 mm coverslips in a 6-well dish and incubated with 10 μM BrdU (Invitrogen) for 12 to 24h. Lipofectamine 2000 (Invitrogen) was used for transfection of 1 μg RNF168-mCherry or SET8-GFP plasmids. Cells were micro irradiated using a PALM MicroBeam Laser Capture microscope (Zeiss) (355nm UV laser). Briefly, the coverslips were transferred to a magnetic incubation chamber (Chamlide) and overlaid with 1 mL medium. Coverslips were mounted on the microscope and irradiated with a pre-determined pattern using 20% cutting speed, 55% focus and 30% power through a 40× objective. The time required to complete the irradiation pattern was approximately 8 minutes. Cells were allowed to recover at 37°C for 30 min before being processed for immunofluorescence. It was empirically determined that these settings did not produce a detectable DNA damage response in unsensitized cells (data not shown).

### Immunofluorescence and microscopy

Cells were seeded on 0.17 mm glass coverslips and treated as indicated in the figure legends. Cells were rinsed 2× with PBS, preextracted with CSK for 5 min at 4 degrees, fixed with 3.7% paraformaldehyde in PBS for 15 min at room temperature and washed 4× with PBS. Cells were permeabilized by incubation in PBS containing 0.25% Triton X-100 (v/v) for 15 min at room temperature and rinsed 1× with PBS. Samples were blocked with antibody dilution buffer (5% BSA, 0.1% Triton X-100 in PBS) for approximately 1h and incubated with the indicated primary antibody overnight at 4 degrees or at room temperature for 2 h. Secondary antibodies were incubated for 45 min at room temperature. Coverslips were mounted with Vectashield + DAPI (Vector Laboratories) and Prolong Diamond containing DAPI. The fluorescence from GFP fusion proteins was imaged from the intrinsic GFP fluorescence remaining after processing. Images were acquired using a Zeiss Axio Observer Z1 microscope using a 40× oil immersion objective and standard FITC, TRITC and Cy5 filter sets.

### Antibodies

We used the following antibodies: mouse-gH2AX (ab81299, Abcam), rabbit anti-SET8 (2996, Cell signaling Technology), rabbit anti-53BP1 (NB100-305, Novus Biologicals), rabbit anti-RNF168 (ab220324, Abcam), rabbit anti-beta actin (ab8227, Abcam).

### RNA interference

RNF8 and RNF168 siRNA’s transfected using Lipofectamine RNAiMAX (Invitrogen) at 100 nM concentration. Sequences of siRNAs used in this work are as follows:

CONTROL: 50-UAAGGCUAUGAAGAGAUAC-3 (Dharmacon),
RNF8si #1: 5’-GGACAAUUAUGGACAACAA-3’ (Dharmacon)
RNF8si #2: 5’-UGCGGAGUAUGAAUAUGAA-3’(Dharmacon)
RNF168 si #1: 5’- GGCGAAGAGCGAUGGAGGATT-3’ (Ambion)
RNF168/si#2: 5’- GACACUUUCUCCACAGAUAUU-3’ (Dharmacon)

### GFP-Trap based Immunoprecipitation (Figure 3A)

Cells were washed twice with ice cold PBS, scraped and spin down for 3 min at 300g. Cells were re-suspended in the Buffer A (20 mM HEPES, 10 mM KCl, 1.5 mM MgCl, 0.34 M Sucrose, 10% Glycerol, 5mM BME). Triton X100 was added to final concentration 0.02%, homogenized by inversion and incubated on ice for 10 minutes by inverting the tube every 2 minutes. Nuclei was pelleted via centrifugation for 1300g for 5 min at 4°C and then re-suspended in the Buffer B (50mM Tris pH 7.5, 500mM NaCl, 1mM EDTA, 0.1% NP-40, 20% Glycerol, 1 mM DTT) with 5 -10 strokes through 19.5 G needle. Benzonase was added and samples were incubated on ice for 30 min by inverting the tube every 5 minutes. Samples were sonicated for 5 min using Bioruptor sonicator at high intensity mode. Nuclear extracts were diluted to final concentration 150 mM NaCl, using No Salt Buffer B (50 mM Tris pH 7.5, 1mM EDTA, 0.1% NP-40, 20% Glycerol, 1mM DTT). After centrifugation (20,000 g, 4°C, 10 min), supernatants were transferred into clean tubes. SDS-PAGE sample buffer was added to 5% - 10% of diluted sample and referred as “input fraction”.

The beads (GFP-Trap agarose) were equilibrated by washing with 150mM NaCl Buffer B. 20 μl GFP-Trap agarose was added into 500 μl lysate and incubated from 10 min to 2h. Immunocomplexes were harvested by centrifugation and supernatants were collected, SDS-PAGE sample buffer was added and aliquoted as “non-bound fraction”.

The beads were washed twice by Buffer 1 (50 mM Tris pH 7.5, 150 mM NaCl, 1 mM EDTA, 0.1% NP-40, 20% Glycerol, 1 mM DTT) and Buffer 2 (50 mM Tris pH 7.5, 0-500 mM NaCl, 1 mM EDTA, 0.1% NP-40, 20% Glycerol, 1 mM DTT). Beads were re-suspended in 50-100 μl SDS-PAGE Sample Buffer and referred as “bound fraction”.

Proteins were eluted by boiling at 95°C for 10 min. For immunoblot analysis 1% of input and e.g. 20% of bound fractions subjected to SDS-PAGE, transferred to a nitrocellulose or PVDF membrane to detect GFP-fusion protein with an α-GFP antibody and interacting proteins with the respective antibodies.

### Chromatin recruitment experiment (Figure 3B)

Hela cells with 256 LacO array and the LacR plasmid were generous gift from Daniel Durocher (1). Cells were transfected with LacR plasmid using Lipofectamine 2000 (Invitrogen) for 48 hrs, micro irradiated and immunostained for SET8 and RNF168.

## Acknowledgments

We thank Dan Durocher for the mCherry-RNF168 plasmids and for the HeLa-lac0 cell line. We thank Jiri Lukas for siRNAs and siRNA-resistant RNF168 plasmids.

